# Genetic redundancy in iron and manganese transport in the metabolically versatile bacterium *Rhodopseudomonas palustris* TIE-1

**DOI:** 10.1101/2020.05.08.085498

**Authors:** Rajesh Singh, Tahina Onina Ranaivoarisoa, Dinesh Gupta, Wei Bai, Arpita Bose

**Author notes:** Corresponding author: Arpita Bose, Department of Biology, Washington University in St. Louis, One Brookings Drive, St. Louis, MO, 63130. These authors contributed equally to this work with author order based on writing contribution (Rajesh Singh, Tahina Onina Ranaivoarisoa).

## Abstract

The purple non-sulfur bacterium *Rhodopseudomonas palustris* TIE-1 can produce useful biochemicals such as bioplastics and biobutanol. Production of such biochemicals requires intracellular electron availability, which is governed by the availability and the transport of essential metals such as iron (Fe). Because of the distinct chemical properties of ferrous [Fe(II)] and ferric iron [Fe(III)], different transport systems are required for their transport and storage in bacteria. Although Fe(III) transport systems are well characterized, we know much less about Fe(II) transport systems except for the FeoAB system. Iron transporters can also import manganese (Mn). Here, we study Fe and Mn transport by five putative Fe transporters in TIE-1 under metal-replete, -deplete, oxic and anoxic conditions. We observe that by overexpressing *feoAB, efeU*, and *nramp1AB*, the intracellular concentration of Fe and Mn can be enhanced in TIE-1, under oxic and anoxic conditions, respectively. The deletion of a single gene/operon does not attenuate Fe or Mn uptake in TIE-1 regardless of the growth conditions used. This indicates that genetically dissimilar yet functionally redundant Fe transporters in TIE-1 can complement each other. Relative gene expression analysis shows that *feoAB* and *efeU* are expressed during Fe and Mn depletion under both oxic and anoxic conditions. The promoters of these transporter genes contain a combination of Fur and Fnr boxes suggesting that their expression is regulated by both Fe and oxygen availability. The findings from this study will help us modulate intracellular Fe and Mn concentration, ultimately improving TIE-1’s ability to produce desirable biomolecules.

**IMPORTANCE:** *Rhodopseudomonas palustris* TIE-1 is a metabolically versatile bacterium that can use various electron donors including Fe(II) and poised electrodes for photoautotrophic growth. TIE-1 can produce useful biomolecules such as biofuels and bioplastics during various growth conditions. Production of such reduced biomolecules is controlled by intracellular electron availability, which in turn is mediated by various iron-containing proteins in the cell. Several putative Fe transporters exist in TIE-1’s genome. Some of these transporters can also transport Mn, part of several important cellular enzymes. Therefore, understanding the ability to transport and respond to varying levels of Fe and Mn under different conditions is important to improve TIE-1’s ability to produce useful biomolecules. Our data suggest that by overexpressing Fe transporter genes via plasmid-based expression, we can increase the import of Fe and Mn in TIE-1. Future work will leverage these data to improve TIE-1 as an attractive microbial chassis and future biotechnological workhorse.

## INTRODUCTION

Iron (Fe) and manganese (Mn) are essential nutrients for biological processes (1-3). Because of the unique redox properties of Fe, it serves an important role as a cofactor in enzymes involved in a variety of cellular processes (4). Similarly, Mn plays an essential role in lipid, protein, and carbohydrate metabolism (5). It also serves as a cofactor of Mn-dependent superoxide dismutase and can contribute to the catalytic detoxification of reactive oxygen species (ROS) (5-7). However, when in excess, these metals are toxic to cells. An evaluation of Fe toxicity revealed that the presence of 1 mM Fe(III) and 0.5 mM Fe(II) can significantly affect the growth of *E. coli* (8). Such elevated concentration can perturb intracellular redox conditions and produce reactive hydroxyl radicals (1). Likewise, excess Mn can inactivate enzymes that use other divalent ions. Mn is known to replace Fe from cellular proteins (6, 7). Therefore, the function of many Fe-containing proteins (e.g., cytochromes, dehydrogenases, and iron-sulfur proteins), which are involved in diverse cellular processes, may be affected due to replacement of Fe by Mn when a very high level of Mn is present (9). The interplay of the need and the toxicity of these metals forces organisms to carefully maintain metal homeostasis. This ensures that metal availability is in accordance with their physiological needs (1). This homeostasis is largely maintained by regulating specific metal transport systems across all biology (4, 10-17).

Depending on oxygen availability in the environment, iron exists in two forms: ferric [Fe(III)] and ferrous [Fe(II)]. In oxic environments, most bacteria acquire insoluble Fe(III) by synthesizing high-affinity Fe(III) siderophores via a process that involves multi-enzyme pathways (1, 18-20). In contrast, Fe(II) uptake is much simpler and involves direct transportation of Fe(II) into the cytoplasm through distinct Fe(II) transporters (21-23). Likewise, depending on oxygen availability, manganese is also found in two most stable forms: Mn(II) and Mn(IV). In many cases, Fe(II) transporters can also transport Mn(II) across the cytoplasmic membrane (24, 25). Although Fe(III) transport systems are well understood, we know much less about Fe(II) transport systems (except for the FeoAB system) and their ability to transport Mn(II) in bacteria, which is the focus of this study.

Here, we study Fe and Mn transport in a metabolically versatile anoxygenic phototroph *Rhodopseudomonas palustris* TIE-1 (TIE-1). Depending on the availability of different energy and carbon sources, TIE-1 can perform multiple modes of metabolisms such as photoautotrophy, photoheterotrophy, chemoautotrophy and chemoheterotrophy (26-28). Similar to the other strains of *Rhodopseudomonas* that have been used to produce value-added compounds (29-34), TIE-1 can also produce compounds such as bioplastics and biofuels under various growth conditions ((35), Bai et al., submitted for publication). Electron transfer during the synthesis of these compounds requires Fe-containing proteins, such as cytochromes. The activity of such proteins depends on the bioavailability and the transport of Fe (28, 35, 36).

We studied the role of five genes *(feoAB, efeU, nramp1AB, nramp3AB*, and *sitABC)* that encode Fe(II) transporters in TIE-1 to import Fe and Mn under metal-replete and -deplete conditions with respect to oxygen availability. Using heterologous complementation, mutant analysis, and gene expression analysis, we show that by manipulating TIE-1’s Fe transport systems, we can enhance its ability to acquire Fe and Mn. Identifying such key transporters might be beneficial for developing TIE-1 as a bioproduction chassis.

## RESULTS

### TIE-1 possesses six putative ferrous iron transporter genes

We identified six putative Fe(II) transporters in TIE-1 using whole-genome homology searches for previously characterized Fe(II) transport systems from *E. coli* and *R. palustris* CGA009 (Fig. 1). The first is the Fe(II)-specific FeoAB system. Unlike *E. coli*, TIE-1 only contains FeoA and FeoB but not FeoC. However, the Feo system lacking FeoC is fully functional for Fe(II) uptake in several bacteria (37). The second Fe(II)-specific transport system is EfeU, which is a member of the oxidase-dependent iron transporters (OFeT) and is a homolog of the iron permease Ftr1p from yeast (38). It is a part of the *efeUOB* tricistronic operon, which is expressed in response to Fe availability in *E. coli* (24). The third system is SitABC (SitD is absent), which has been reported to have a much greater affinity for Mn than Fe (39). Fe and Mn-dependent regulation of *sitABC* operon have been previously described in *Staphylococcus epidermidis* (40) and *S. aureus* (41), respectively. The fourth transport system belongs to the natural resistance-associated macrophage protein (NRAMP) family. We found three *nramp* gene/operons: *nramp1AB, nramp2, and nramp3AB*, which are also known to be important for Fe and Mn uptake and homeostasis in other organisms (42, 43). Nramp1A belongs to the DNA binding transcriptional regulator LysR family whereas Nramp3A is a hypothetical protein (Rpal_3713) that is only predicted in the genome of TIE-1 and the related strain CGA009 (Fig. 1E). Nramp1B, Nramp2, and Nramp3B belong to a Mn transporter protein family, MntH from *E. coli* (42, 43). Because we were unable to delete *nramp2* from TIE-1, we exclude *nramp2* from our studies henceforth.

**Figure 1.**
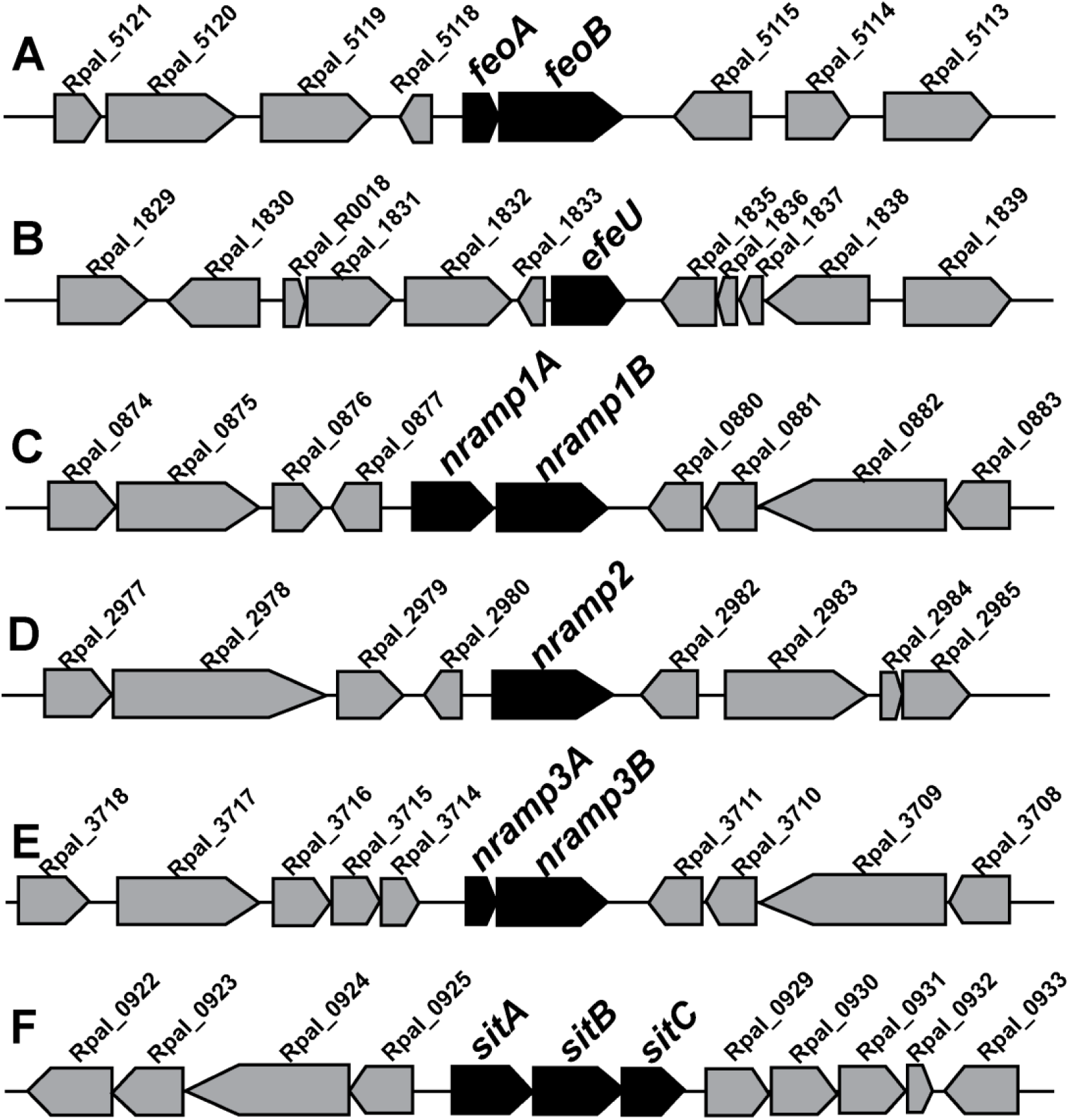
Genomic organization of *R. palustris* TIE-1 genes that are likely involved in Fe(II) transport. Six candidates’ loci were identified (Panel A-F). The genes of interest are shown in black with their names bolded. Other genes are shown in grey with their locus tags searchable at https://img.jgi.doe.gov/cgi-bin/w/main.cgi.

### Putative ferrous iron transport systems from TIE-1 can rescue iron transport deficiencies in a heterologous host

To assess whether the putative Fe(II) transporters support iron uptake, we performed two heterologous complementation experiments in *E. coli* as an initial test. For the first complementation experiment, we used *E. coli* H1771, a mutant defective for the Feo system and siderophore biosynthesis with a chromosomal *fhuF-lacZ* (encoding β-galactosidase) reporter system. This reporter system has been previously used in similar studies (44-46). The *fhuF* promoter of the reporter system is responsive to Fe, where repression of β-galactosidase activity represents adequate intracellular Fe concentration while an increase in β-galactosidase activity indicates Fe starvation (47, 48).

The identified Fe transporter genes/operons from TIE-1 were cloned into the low copy number vector pWKS30 (47) and were driven by the constitutive promoter *P*_*aphII*_ (49). We have also recently reported gene expression using *P*_*aphII*_ promoter in TIE-1 where we produced PioA in TIE-1 in a *pioA*-deleted TIE-1 background using this promoter and confirmed protein expression by Western blot (36). To further confirm the efficiency of the P_*aphII*_ promoter, we built a plasmid where *mCherry* was driven by *P*_*aphII*_. TIE-1 carrying this plasmid showed a five thousand-fold higher fluorescence signal (normalized to OD_600_) compared to wild type TIE-1 (Fig. S1). This promoter has been also used previously in a closely related organism such as *Rhodobacter capsulatus* (50).

Plasmids carrying the transporter genes were then transformed into *E. coli* H1771, and β-galactosidase (reporter) activity was assayed under four different conditions namely, **(i)** Fe(III)-replete [50 µM Fe(III)-citrate]; **(ii)** Fe(III)-deplete [50 µM Fe(III)-citrate with 100 µM DiP (2,2’-dipyridyl)]; **(iii)** Fe(II)-replete [50 µM ascorbate-reduced Fe(III)-citrate]; and **(iv)** Fe(II)-deplete [50 µM ascorbate-reduced Fe(III)-citrate with 100 µM DiP]. The membrane-permeant chelator DiP was used to control the bioavailability of Fe(II) and Fe(III). DiP is a Fe(II)-specific bidentate organic ligand that acts by directly depleting intracellular stores of ferrous iron (51). The inherent *fhuF-laZ* reporter system of *E. coli* H1771 carrying an empty vector expressed *lacZ* to a full extent due to a low intracellular Fe concentration (Fig. 2). *E. coli* H1771 complemented with K12 *feoABC* through a plasmid served as a positive control and showed more than 80% decrease in *lacZ* expression as determined by β-galactosidase activity compared to the vector control under all conditions (Fig. 2A-D). Regardless of different Fe(III) concentrations and transporter genes from TIE-1, none of the complemented *E. coli* H1771 strains showed significant changes in β-galactosidase activity (Fig. 2A-D). In contrast, all the complemented strains under Fe(II)-replete conditions showed a lower β-galactosidase activity (Fig. 2C), compared to both the vector control and the activities under Fe(II)-deplete conditions (Fig. 2D). The most pronounced difference in the β-galactosidase activity compared to the empty vector was observed for the *efeU*-complemented strain under Fe(II)-replete conditions (62% lower than the vector control) (Fig. 2C).

**Figure 2.**
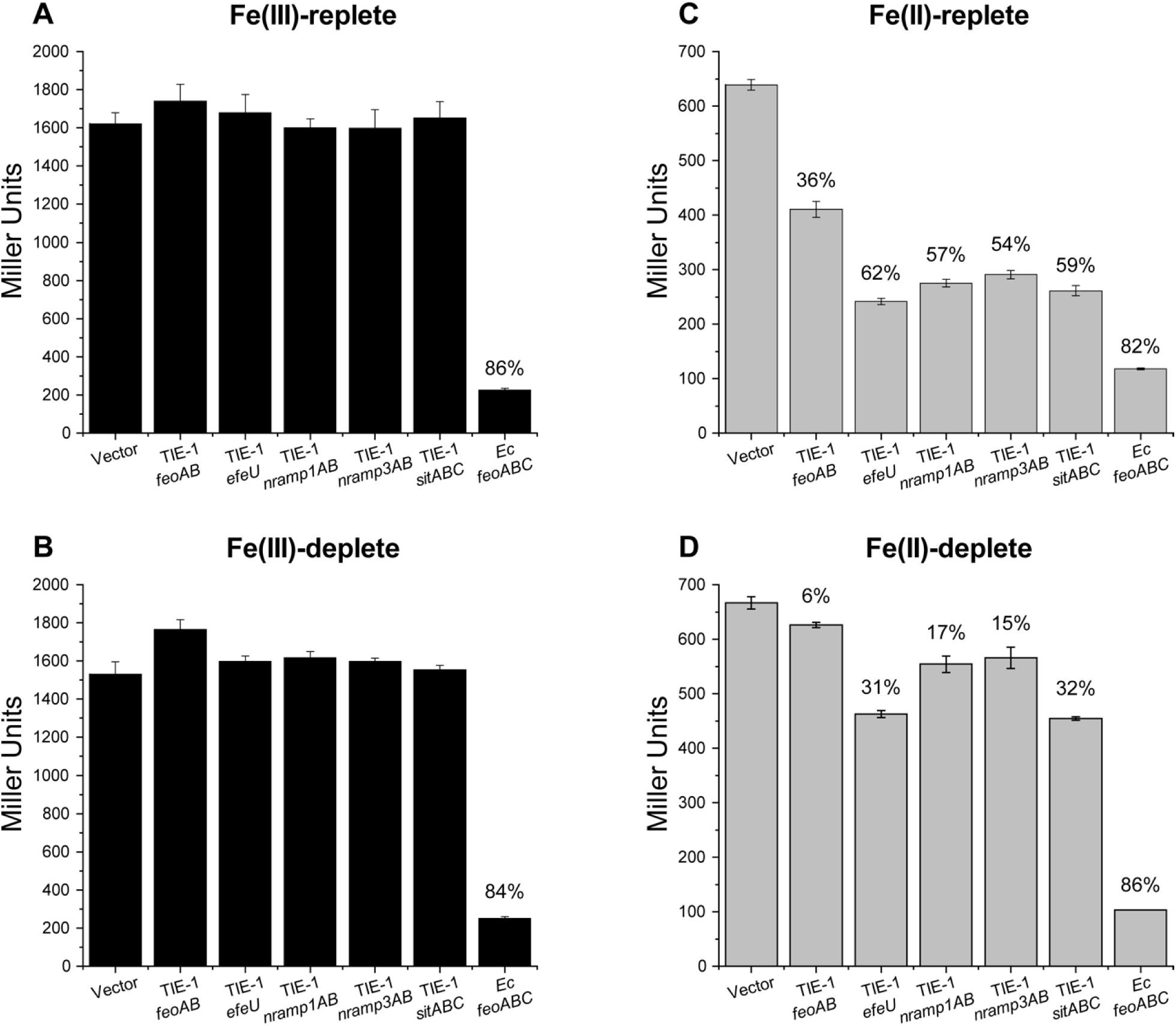
*E. coli* H1771 complemented by selected iron transport genes resulted in less iron stress when grown in Fe(II) supplied LB medium but had no effect when grown in Fe(III) supplied LB medium. The Miller β-galactosidase assay was used to determine the level of iron stress. Four conditions were used: (A) Fe(III)-replete [50 µM Fe(III)-citrate]; (B) Fe(III)-deplete [50 µM Fe(III)-citrate with 100 µM DiP]; (C) Fe(II)-replete [50 µM ascorbate-reduced Fe(III)-citrate]; and (D) Fe(II)-deplete [50 µM ascorbate-reduced Fe(III)-citrate with 100 µM DiP]. Cells at the exponential phase were used for all assays. Values are represented as mean ± standard error of six independent biological measurements. Number on each bar represents the percent reduction (normalized to empty vector control) of Miller β-galactosidase activity. X-axis shows the genes used for complementation. TIE-1 – *Rhosdopseudomonas palustris* TIE-1, Ec – *E. coli* K12.

To further confirm the role of the putative Fe(II) transporters in cell growth, we employed a second heterologous complementation experiment using *E. coli* GR536. *E. coli* GR536 is a mutant lacking all five known Fe transport systems (*ΔfecABCDE::kan ΔzupT::cat, ΔmntH, ΔfeoABC* and *ΔentC)* and, therefore, it is unable to grow under Fe-deplete condition (52). We expressed the Fe(II) transporter genes from TIE-1 in this heterologous host and examined if their expression can rescue the growth defect of *E. coli* GR536 under Fe-deplete conditions (Fig. 3A-E). Indeed, complementation of *E. coli* GR536 with TIE-1’s *efeU, nramp1AB, nramp3AB, sitABC* restored its growth to varying degrees after ∼60 hours (Fig. 3B-E). In contrast, although the growth defect was rescued by complementation with *E. coli* K12’s *feoABC* (a positive control) within 24 hours (Fig. 3F), the expression of TIE-1’s FeoAB system could not rescue the growth defect (Fig. 3A). This result suggests that all three genes of the Feo system including FeoC are important for Fe transport in *E. coli*.

**Figure 3.**
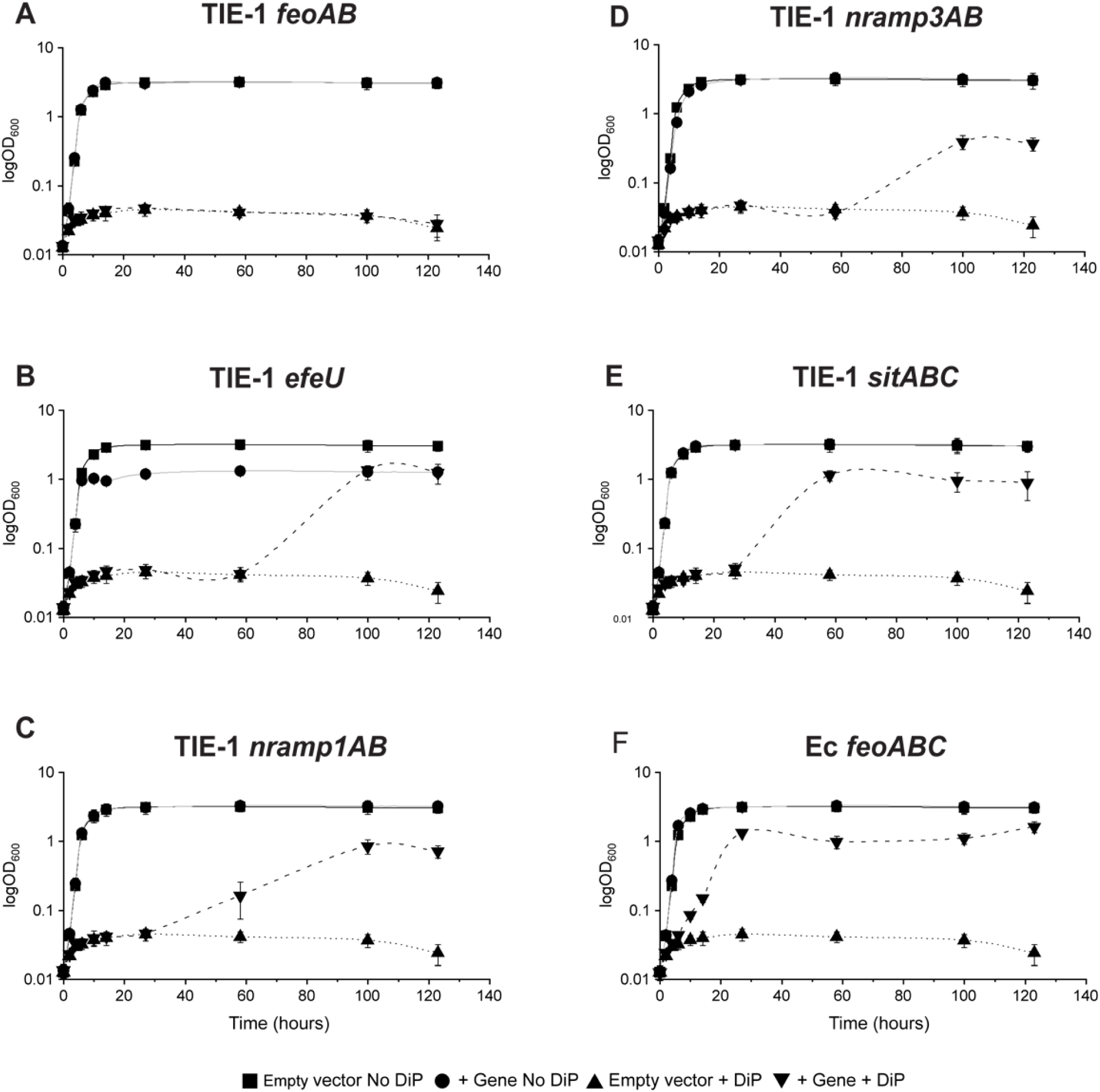
Complementation with selected Fe(II) transport genes aid growth of *E. coli* GR536 mutant (which lacks all known iron transport genes) during severe iron deprivation (in Tris-mineral salts medium at pH 7.0 with no added iron). *E. coli* GR536 was complemented by *Rhodopseudomonas palustris* TIE-1 (A) TIE-1 *feo*AB; (B) TIE-1 *efeU*; (C) TIE-1 *nramp1AB;* (D) TIE-1 *nramp3AB*; (E) TIE-1 *sitABC*; *E. coli* K12 *feoABC* (positive control). Two representative growth conditions are shown: without and with 150 M 2,2’-dipyridyl (DiP). Empty vector was used as control for both conditions. Growth was measured as OD_600_. Values represent the mean ± standard deviation of biological triplicates. TIE-1 – *Rhodopseudomonas. palustris* TIE-1, *Ec* – *E. coli* K12.

### Putative ferrous iron transporters from TIE-1 do not transport zinc, cadmium or copper

Based on previous studies, the FeoAB and EfeU transporters are predicted to be specific for Fe uptake while the Nramp and Sit systems are known to also transport other divalent cations (39, 42, 53, 54). Therefore, we further investigated the metal specificity of these putative Fe transporters in TIE-1. Although DiP is a strong intracellular Fe(II) chelator, it also binds to other divalent metals such as copper [Cu(II)], cadmium [Cd(II)], and zinc [Zn(II)] (55). Because of this broader metal binding capacity of DiP, we further used this chelator in studying the metal specificity of these transport systems. Heterologous complementation was performed in two different strains: **(i)** *E. coli* GG48 which lacks key Zn transport genes, and **(ii)** *E. coli* (Δ*copA* Δ*cueO* Δ*cusCFBA::cat*), which lacks the key Cu transport genes (56). Complementation of these *E. coli* hosts with a functional Zn/Cu/Cd transport system is detrimental and is reflected by a lower growth rate when these metals are provided in the growth medium (Table 1). We then determined the ability of the putative Fe(II) transport systems from TIE-1 to transport Zn(II), Cd(II), and Cu(II) by monitoring the growth rate of the heterologous *E. coli* host complemented with plasmids expressing TIE-1’s putative Fe(II) transporters compared to the control strain (Table 1, *P*≤0.05). These data suggest that the TIE-1 Fe(II) transporters had either no effect or a positive effect on the growth rate of *E. coli*. The increase in growth rate could be accounted for the ability of these transporters to bring in extracellular Fe into the cell. The only case where we observed a negative effect on growth rate was under the Cd/Cu-supplemented conditions for the *E. coli* (Δ*copA* Δ*cueO* Δ*cusCFBA::cat*) complemented with *E. coli feoABC* system. Overall, these data suggest that the TIE-1’s Fe transport systems do not import enough Zn/Cu/Cd into these *E. coli* mutant strains to observe a severe growth defect. The result implies that these TIE-1 Fe transport systems likely have a low affinity for Zn/Cu/Cd.

**Table 1.** Metal specificity tests using *E. coli* GG48 for Zn and *E. coli* (Δ*copA* Δ*cueO* Δ*cusCFBA::cat*) for Cd and Cu transformed with plasmids with genes encoding Fe transporters from *R. palustris* TIE-1 (*Rp*) or *E. coli* K12 (*Ec*).

| Gene/Operon | Zn | Cd | Cu |
| --- | --- | --- | --- |
| *Rp feoAB* | 0 | + | 0 |
| *Rp efeU* | 0 | 0 | 0 |
| *Rp nramp1AB* | + | 0 | 0 |
| *Rp nramp3AB* | 0 | + | 0 |
| *Rp sitABC* | 0 | + | 0 |
| *Ec feoABC* | + | - | - |

A “0” indicates no significant difference in toxicity compared to the empty vector control ( $P>0.05$ ), “+” a significantly higher growth rate in the experimental strain versus control, and “-” a significantly lower growth rate versus control ( $P\leq 0.05$ ).

### Overexpression of ferrous iron transporter genes increases intracellular iron and manganese in TIE-1 under oxic and anoxic conditions

After confirming the role of the putative Fe(II) transporters in Fe transport and growth of the heterologous host *E. coli*, we sought to investigate their role in Fe and Mn accumulation in the native organism, TIE-1. We constructed markerless deletions of these genes in TIE-1 as described in the method section (27). We were unable to obtain some of the mutants despite screening >100 colonies. These include *feoA, feoB, nramp1A, nramp2*, and *sitA*. We tested the ability of the five obtained mutants *(ΔfeoAB, ΔefeU, Δnramp1AB, Δnramp3AB, ΔsitABC)*, their complemented strains (each complemented by corresponding genes) and WT (positive control) to intracellularly accumulate Fe and Mn under anoxic, oxic, metal-replete and -deplete conditions. The complemented transporter genes were expressed using the same plasmid, from the same *P*_*aphII*_ promoter and ribosome binding site described above. Therefore, we assume that the expression level for each gene is similar. We quantified intracellularly accumulated Fe and Mn using inductively coupled plasma mass spectrometry (ICP-MS). The results are expressed as the ratio of metal accumulated by a strain normalized to the metal accumulated by the wild-type control strain. We did not observe any metal precipitation during our experiments.

TIE-1 is an ideal candidate to study Fe and Mn uptake due to its ability to grow under both oxic and anoxic conditions that allows us to study Fe(II), Fe(III), Mn(II) and Mn(IV) transport systems using the same organism. Although the ideal conditions to study the transport of these metal species is to use the same growth medium, we had to grow TIE-1 aerobically on yeast peptone (YP) medium and anaerobically on defined freshwater (FW) medium as described in the methods section. This is because TIE-1 does not grow robustly in YP medium under anaerobic conditions, or in FW medium under aerobic conditions. Such different growth conditions have recently been used in a Fe transport study in *Shewanella oneidensis* (57). To further control the bioavailability of Fe and Mn, two different chelators EDTA and DiP were tested for all conditions. Being membrane-permeant (58), DiP has the potential to diffuse into cells and alter the intracellular metal distribution (59). It has a stability constant of 17.5 and 16.3 with Fe(II) and Fe(III), respectively and 6 with Mn(II) (60, 61). In contrast, EDTA is a large molecule that is membrane-impermeant (58). In addition, the presence of two nitrogen atoms and four carboxyl groups in EDTA’s structure makes it a strong metal chelator for iron [stability constant 25.10 with Fe(III) and 14.33 with Fe(II)] and manganese [stability constant 24.80 with Mn(III) and 14.04 with Mn(II)] (59). Due to its inability to enter cells, it can only bind to extracellular Fe or Mn via metal-ligand interaction eventually limiting their transports (58). Our results indicate that the complemented strains accumulate comparable amounts of Fe as the WT and the mutant strains under anoxic growth condition. When Fe was depleted by adding EDTA or DiP under anoxic condition, growth was suppressed (OD_660_∼0.1, Fig. S2) and Fe acquisition was abolished (Fig. 4A-E). Two exceptions were Δ*nramp3AB* and Δ*sitABC* where growth was observed with EDTA and/or DiP in the mutants but not in the WT or the complemented mutants (Fig. S2). Because we could not obtain measurable metal concentration from the WT with which we normalized the other values, we could not report the Fe concentration from Δ*nramp3AB* and Δ*sitABC* during growth with EDTA and/or DiP.

**Figure 4.**
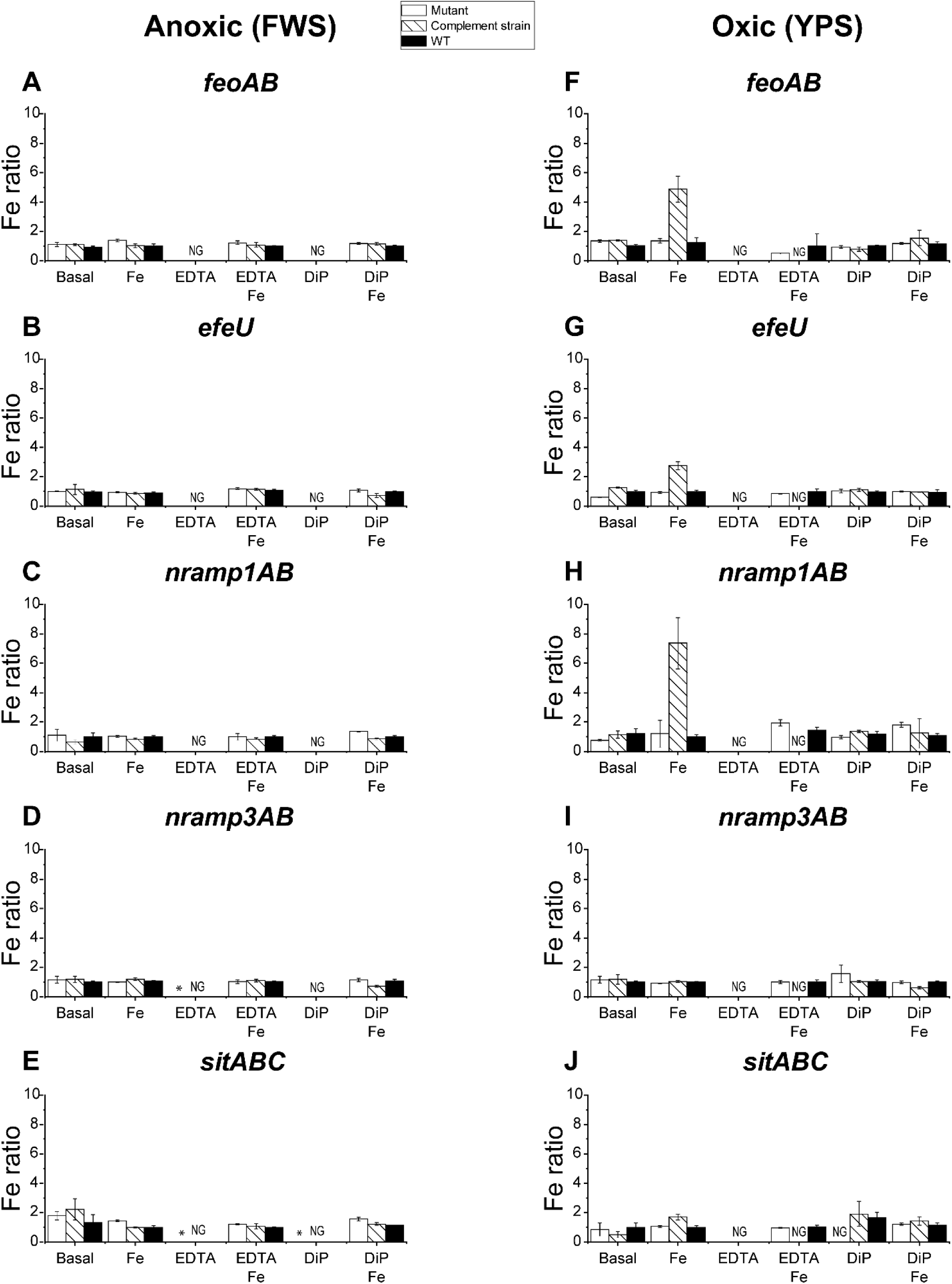
Fe acquisition by mutants, complemented strains and *Rhodopseudomonas. palustris* TIE-1 (WT) under Fe-replete, -deplete, oxic and anoxic conditions using inductively coupled plasma mass spectrometry (ICP-MS). The iron content of the cell is normalized to the Fe acquisition by WT. (A-E) Fe acquisition under anoxic conditions. (F-J) Fe acquisition under oxic conditions. Values represent the mean ± standard deviation of biological triplicates. DiP-2, 2’-dipyridyl; EDTA-ethylenediaminetetraacetic acid; FWS – freshwater media supplemented with 1 mM succinate; YPS – yeast peptone supplemented with 1 mM succinate. NG- No growth. * – these conditions were where OD_660_ >0.1 was observed only for the mutant but not WT or the complemented strain (see Fig. S2).

Supplementation of 50 µM Fe(II) did not increase Fe accumulation by these strains under anoxic condition (Fig. 4A-E; Table S1 a-e). In contrast, compared to the WT and their respective mutants, Fe(II) supplementation to the basal YP (yeast peptone supplemented with 1 mM succinate) medium under oxic condition significantly increased intracellular Fe concentration (up to ∼2 to ∼7-fold higher) by the complemented strains of *ΔfeoAB, ΔefeU, Δnramp1AB*, and *ΔsitABC* (Fig. 4F-H, *P*<0.05, Table S1). Similar to the anoxic growth condition, the addition of EDTA suppressed cell growth (OD_660_<0.1) as well as Fe transport by all the strains (Fig. 4F-J). However, Fe(II) supplementation restored Fe transport and rescued their growth except for the complemented strains. Because gentamicin was added to maintain plasmid [pBBRIMCS-5] (62) expressing the individual genes from TIE-1 in the complementation strains, we hypothesize that the lack of growth is likely because of a combined effect of gentamicin and EDTA. A combination of EDTA and gentamicin has been previously reported to inhibit the growth of *Pseudomonas aeruginosa* (63). The growth inhibition in the presence of EDTA was consistently observed in all the complemented strains under oxic conditions despite the addition of Fe (Fig. S2k). In contrast, the same effect did not hold true under anoxic conditions with freshwater (FW) medium. Even the complemented strains accumulated a very similar amount of Fe compared to the mutants and WT strains with their subsequent growth (Fig. 4A-E, Fig. S2). The combined toxic effect of EDTA and gentamicin might have been reversed by the presence of other cations in the FW medium including magnesium and calcium. The addition of magnesium, calcium and iron appears to block cell death and detachment of *P. aeruginosa* biofilms (63). This could suggest that the addition of other divalent metals increases the binding competition to EDTA and frees more Fe for cell growth. However, future work will be required to determine the specific role of medium composition in this process. In contrast, combination of DiP and gentamicin under oxic condition does not seem to have any toxic effects to the complemented strains and their subsequent metal accumulation (Fig. 4F-J).

To investigate if the putative Fe(II) transporters also play a role in Mn transport, we quantified the total amount of Mn accumulated by the different strains of TIE-1. Under anoxic growth conditions with a supply of Mn and EDTA, the complemented strains of *ΔfeoAB* and *ΔefeU* accumulated noticeably higher Mn (∼3.6 and ∼1.8-fold, respectively) compared to WT and their mutants (Fig. 5A and B, *P*<0.05, Table S1 k and l). However, when the strains were cultured in the basal FW medium supplemented with Mn(II), only the *ΔefeU* complemented strain accumulated a significant amount of Mn compared to WT and its mutant (Fig. 5B, Table S1 l). Similar to the Fe studies, the addition of EDTA and DiP to the basal medium abolished cell growth and Mn accumulation by all the strains under anoxic condition with the two exceptions noted above (Fig. 5A-E, Fig. S2).

**Figure 5.**
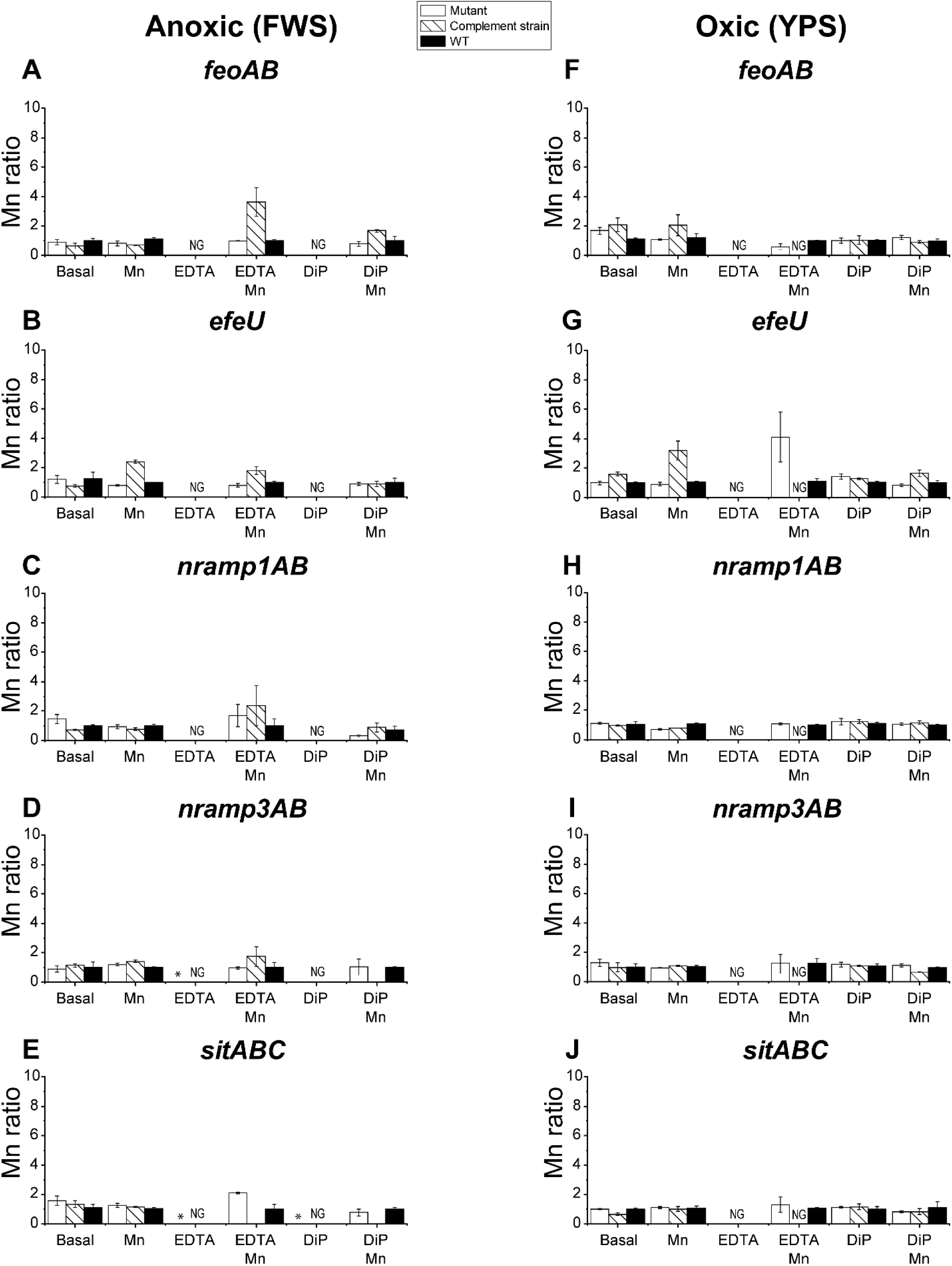
Mn acquisition by mutants, complemented strains and *R. palustris* TIE-1 WT under Mn-replete, -deplete, oxic and anoxic conditions. The Mn content of the cell is normalized to the Mn acquisition by WT. (A-E) Mn acquisition under the anoxic conditions. (F-J) Mn acquisition under oxic conditions. Values represent the mean ± standard deviation of biological triplicates. DiP-2, 2’-dipyridyl; EDTA-ethylenediaminetetraacetic acid; FWS – freshwater media supplemented with 1 mM succinate; YPS – yeast peptone supplemented with 1 mM succinate. NG-No growth.* – these conditions were where OD_660_ >0.1 was observed only for the mutant but not WT or the complemented strain (see Fig. S2).

Under oxic growth condition, only the *ΔefeU* complemented strain accumulated a higher amount of Mn compared to the WT as well as the mutant (∼3-fold) when Mn(II) was added to the basal YP medium (Fig 5G, *P*<0.005, Table S1 q). Mn accumulation by the rest of the TIE-1 strains was similar under oxic condition (Fig. 5F-J, Table S1 p-t). The combined toxic effect of EDTA and gentamicin under oxic conditions was also observed here. None of the complemented strains accumulated Mn, when the medium was supplemented with Mn(II) and EDTA, (Fig. 5F-J, Fig. S2 f-j) including the complemented strains of Δ*nramp1AB*, Δ*nramp3AB*, and Δ*sitABC* which showed modest growth (OD_660_∼0.1) (Fig. S2, Fig. h-j). The amount of Mn accumulated by these three strains was below our detection limit. Under Mn-deplete condition using EDTA, we observed neither growth (OD_660_<0.1) nor Mn accumulation by these strains (Fig. 5F-J, Fig. S2). Overall, the Mn accumulation results show that the Fe transporter *efeU* also participates in Mn transport in TIE-1 (Fig. 5G).

### Gene expression of ferrous iron transporters varies in response to iron and manganese availability and oxygen tension

Because Fe transporter genes are regulated primarily by the availability of Fe and oxygen in the environment (64, 65), we analyzed the gene expression profile of these transport systems in wild-type TIE-1 in response to Fe, Mn, and oxygen availability. We used reverse transcription quantitative polymerase chain reaction (RT-qPCR) with *recA* and *clpX* as reference genes as performed previously (28). Data reported here are with respect to *recA* for consistency with previous studies (28, 35, 36, 64, 65). The trends obtained were similar to *clpX* as a reference gene. Under conditions where no growth was observed (OD_660_<0.1), we collected large volumes of cells to extract sufficient RNA for RT-qPCR.

Fe depletion by the addition of EDTA elevated the expression of *feoB* (up to 20-fold), *efeU* (2.5-fold), *nramp3B* (5-fold) and *sitA* (5.6-fold) under anoxic condition (Fig. 6A-E and Table S2 a-e). This overexpression was reversed when the media was supplemented with 50 µM Fe(II). Similarly, Fe depletion by DiP increased the expression of *feoB* by ∼8-fold. However, this upregulation was not reversed by Fe(II) supplementation (Fig. 6A, *P*<0.05, Table S2 a). Among these genes, *nramp1B* appears to be less responsive to Fe bioavailability as the addition of chelators or Fe did not show a significant effect on its expression (Fig. 6C, Table S2 c).

**Figure 6.**
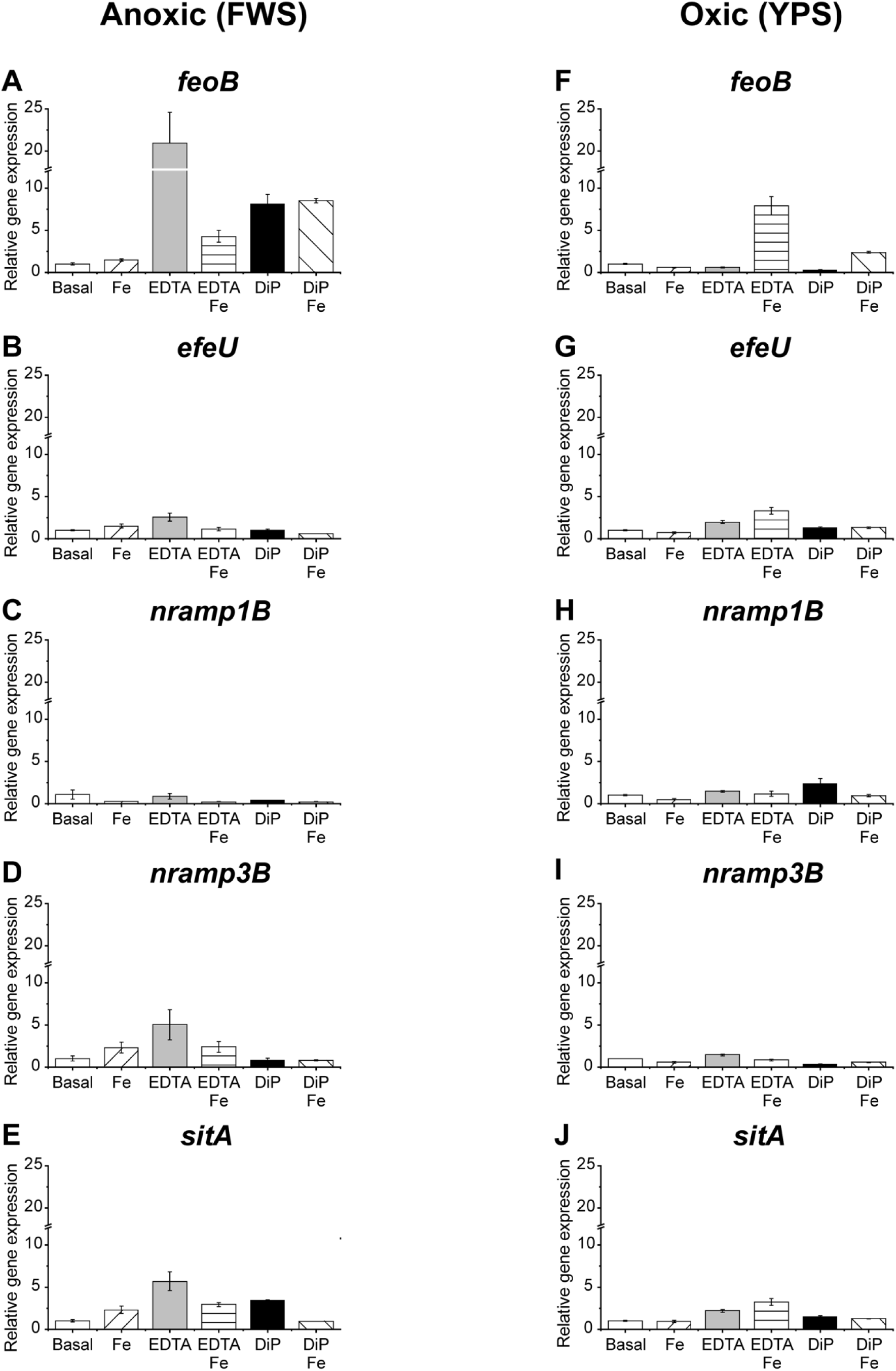
Determination of relative mRNA abundance of Fe transporting genes in TIE-1 using RT-qPCR under Fe-replete, -deplete, oxic and anoxic conditions. (A-E) Relative gene expression under anoxic conditions. (F-J) Relative gene expression under oxic conditions. Values represent the mean ± standard deviation of biological triplicates. DiP-2, 2’-dipyridyl; EDTA-ethylenediaminetetraacetic acid; FWS – freshwater media supplemented with 1 mM succinate; YPS – yeast peptone supplemented with 1 mM succinate.

Under oxic growth condition, the addition of both Fe(II) and EDTA increased the expression of *feoB* (∼8-fold), *efeU* (∼3-fold) and *sitA* (∼3-fold) genes (Fig. 6F, G, J; *P*<0.05, Table S2 f, g, and j). Overall, relative gene expression obtained in the presence of EDTA was higher compared to the expression observed in the presence of DiP (Fig. 6F-J). *nramp3B* was minimally expressed (Fig. 6I).

We also tested the relative gene expressions under different Mn availability. Under anoxic Mn-deplete condition (with EDTA), all five genes namely *feoB, efeU, nramp1B, nramp3B, and sitA* were upregulated (up to ∼2.7-fold) (Fig. 7A-E, Table S2 k-o). However, the expression was downregulated when excess Mn(II) was added. This pattern remained consistent even when the chelator was changed from EDTA to DiP, except for the *feoB* gene (Fig. 7A). Here, the supply of Mn(II) upregulated *feoB*, similar to the trend observed with Fe(II). Unlike the highly variable gene expression observed under anoxic conditions, gene expression under oxic conditions did not vary significantly compared to the control condition (Fig. 7 F-J, Table S2 p-t). We observed that expression of the previously known Fe-specific transporter systems such as Feo and EfeU can be regulated by Mn bioavailability in TIE-1.

**Figure 7.**
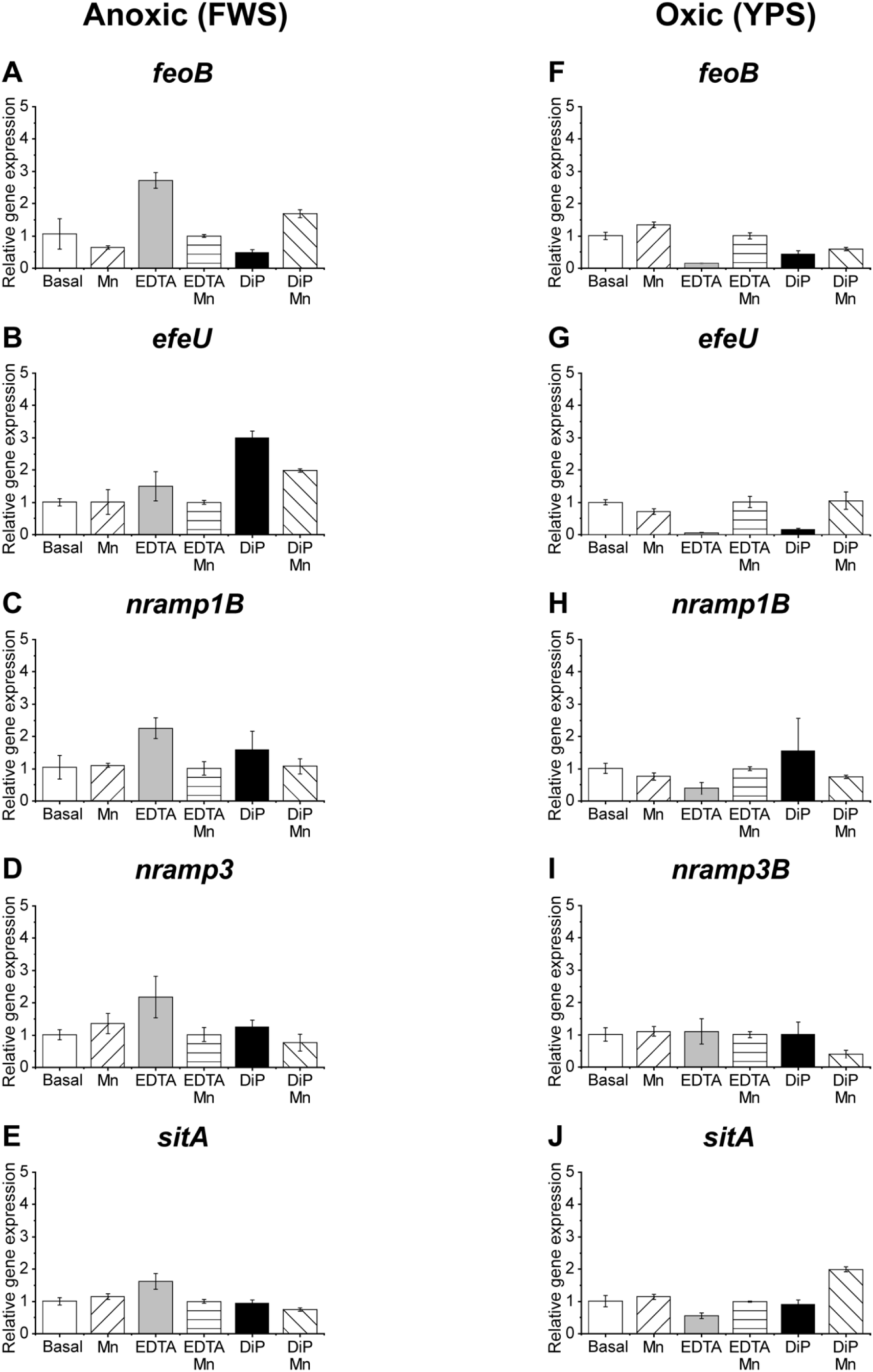
Determination of relative abundance of Fe transporting genes in TIE-1 using RT-qPCR under Mn-replete, -deplete, oxic and anoxic conditions. (A-E) Relative gene expression under anoxic conditions. (F-J) Relative gene expression under oxic conditions. Values represent the mean ± standard deviation of biological triplicates. DiP-2, 2’-dipyridyl; EDTA-ethylenediaminetetraacetic acid; FWS – freshwater media supplemented with 1 mM succinate; YPS – yeast peptone supplemented with 1 mM succinate.

### Promoter predictions

Because Fe transporter genes are regulated primarily by the availability of Fe and oxygen in the environment, next we analyzed the promoter regions of these Fe transporters to identify the binding sites for Fe and oxygen responsive regulators. We found a putative Fur-like binding site (Table S3). Fur is known to be a transcriptional regulator of Fe transporter genes in *E. coli* and many other bacteria (66, 67). We also identified a putative binding site for Fnr family proteins in the *feoAB* promoter along with a Fur/Irr binding site (Fig. S3) (68). Fnr is a transcriptional activator under anoxic condition (69) that turns on the transcription of the *feo* operon in the absence of oxygen (22). The *efeU* promoter has a putative Fnr-box (Fig. S4) but only a weak Fur-box (68). Similar binding sites are identified in the Nramp1AB promoter sequence (Fig. S5). We found a weak Fur-box and an Fnr-box in the *nramp3B* upstream region that encompasses the small ORF *nramp3A*. We were unable to detect *nramp3A* mRNA under any condition tested, which suggests that this ORF is likely not transcribed. The *sitABC* promoter also contains an Fnr box along with a Fur box (Fig. S7) (68).

## DISCUSSION

This study provides insight into the role of five putative Fe(II) transporter genes (Fig. 1) to transport Fe and Mn in TIE-1 under differing Fe, Mn, and oxygen availability. The important role of these transporters in Fe uptake and cellular growth was determined first by heterologous complementation of *E. coli* H1771 (lacking Feo system and siderophores biosynthesis genes) and *E. coli* GR536 (lacking all Fe transporter systems), respectively (Fig. 2D, Fig. 3). Total Fe and Mn quantification in the wild type, single-gene/operon deletion mutants and complemented strains of *R. palustris* TIE-1 showed that these transporter genes are important for Fe and Mn transport in TIE-1. Under anoxic condition, all TIE-1 mutant strains accumulated similar amounts of Fe suggesting that the deletion of no single transporter gene affects iron homeostasis in TIE-1. However, some TIE-1 strains under oxic conditions, particularly, the complemented strains of *feoAB, efeU, nramp1AB* and *sitABC* accumulated a higher amount of Fe compared to WT and mutants (Fig. 4F-J). Under oxic conditions, Fe likely exists as Fe(III). Because overexpressing Fe(II) transporters increases total Fe content under oxic conditions, it is possible that assimilatory ferric reductases may allow reduction of Fe(III) to Fe(II). Fe(II) in the periplasm is subsequently transported by these Fe(II) transporters (present in the inner membrane). This would lead to an increase in intracellular Fe concentration when these transporters are overexpressed under a strong constitutive promoter under oxic conditions. Ferric reductases are known to be involved in Fe acquisition under oxic conditions in several microbes (70). We also found that expressing the putative Fe(II) transporters including the previously described Fe-specific transporters such as *feoAB* and *efeU* can increase intracellular Mn in TIE-1 under both anoxic and oxic conditions (Fig. 5). Under oxic conditions, Mn likely exists as Mn(IV). Similar to Fe(III) uptake, Mn reductases likely mediate Mn(IV) reduction to Mn(II) in the periplasm (71) which can be then transported via Fe transporters.

Our metal accumulation data are also supported by the relative gene expression of these transporters under anoxic and oxic conditions with respect to metal availability. Production of metal transporters are often induced as the metal concentrations are lowered (72, 73). Because the presence of metal chelators reduces metal availability, we observed the upregulation of putative Fe transporters under anoxic metal-deplete conditions (Fig. 6A-E; Fig. 7A-E). A similar result has been observed previously in *Salmonella enterica* where the FeoB protein was overexpressed when levels of both oxygen and Fe were low (74). However, a supply of Fe(II) significantly downregulated the expression. This trend appears more obvious in *feoB* with iron. Upstream regulatory element such as ferric iron regulator *(fur)* has been widely found to regulate the expression of *feo* in the presence or absence of iron (22, 75). When iron is limited by chelators, Fur releases repression upregulating the expression of iron transport genes (22, 75). Fur protein has been also indicated to directly or indirectly mediate manganese-dependent regulation of gene expression (76). Likewise, fumarate and nitrate reductase transcriptional regulator (Fnr) which is a transcriptional activator of anaerobic respiratory genes (69) is known to activate transcription of Feo system under anoxic conditions (48) that can increase the expression level of Feo up to three-fold (22). Indeed, the genome of TIE-1 appears to have at least 11 identifiable Fnr family proteins (Table S3) that are involved in oxygen sensing (50).

Curiously, we observed opposite gene expression trends under oxic conditions. Here, the expression of *feoB and efeU* in the basal YP medium was lower with EDTA compared to the expression obtained in the control conditions (Fig. 6 and 7). We also observed growth inhibition of WT strains with EDTA (Fig. S2 f-j). EDTA has been reported to kill both planktonic cells and biofilms of *P. aeruginosa* (63). We speculate that EDTA may have prevented TIE-1’s growth (Fig. S2 f-j) and subsequent gene expression of *feoB* and *efeU*. This could be also true in the case of the growth with Mn (Fig. 7 F-J). However, the addition of Fe or Mn may reduce the negative effect of EDTA via its interaction with added metals. This is also reflected by their increased growth when Fe or Mn was added to the YP medium (Fig. S2 f-j). Therefore, growth and metal limitation in the presence of Fe and EDTA may have contributed to the increased relative gene expression of *feoB* and *efeU* under oxic condition. However, under anoxic condition, the presence of other divalent cations such as Ca and Mg in FW medium (stability constants of 10.6 and 8.7, respectively) (77) may increase the binding competition to EDTA freeing more Fe for cells. Future work, however, will be required to confirm this hypothesis.

Overall, our results show that TIE-1 tightly regulates and maintains Fe and Mn homeostasis. The deletion of any single gene or operon does not affect the metal uptake capacity and viability of TIE-1. This suggests that the existence of compensatory systems for metal uptake in TIE-1 is crucial for its metabolic adaptability under different levels of Fe/Mn and oxygen.

*R. palustris* TIE-1 has homologs for both Fur and two Irr proteins that mediate iron-dependent regulation of gene expression (Table S3) (68). The genome of TIE-1 appears to have at least 11 identifiable Fnr family proteins (Table S3) that are involved in oxygen sensing (78). One of these homologs is the FixK protein that has a Fe-S cluster and its oxygen sensing response is linked to the two-component regulation system FixLJ (64).

We have recently shown that TIE-1 can produce bioplastics and biofuels such as *n*-butanol under various growth conditions ((35), Bai et al., submitted for publication). Other strains of *Rhodopseudomonas* have also been used to produce value-added compounds (29-34). Production of such biomolecules is controlled by intracellular electron availability, which in turn is modulated by the action of various Fe-containing proteins. Mn is part of many enzymes in central metabolism. By controlling Fe and Mn intake via synthetic biology, we can modulate bioproduction by *R. palustris* TIE-1. This will be investigated in future studies where overexpression of the two Fe transporters *feoAB* and *efeU* (involved in Fe and Mn transport that we have identified here) will be performed in bioplastic and biofuel production strains to see if product titers can be increased.

## EXPERIMENTAL PROCEDURES

### Bacterial strains and growth conditions

*E. coli* strains were obtained from the sources indicated in Table S4. They were grown in Lysogeny Broth (LB) at pH 7.0 at 37 °C with shaking at 225 rpm. *Rhodopseudomonas palustris* TIE-1 was obtained from Dianne K. Newman, Division of Biology and Biological Engineering and the Division of Geological and Planetary Sciences, California Institute of Technology. Prior to photoferrotrophy [Fe(II) as an electron donor], TIE-1 cells were grown chemoheterotrophically in 0.3% yeast extract and 0.3% peptone (YP) medium, with 1 mM sodium succinate and 10 mM MOPS [3-N (morpholino) propanesulfonic acid] at pH 7.0 in the dark at 30 °C with shaking at 250 rpm (64). Time-course cell growth was monitored using Spectronic 200 (Thermo Fisher Scientific, USA). To adapt the cells to photoautotrophic growth using H_2_ as the sole electron donor, chemoheterotrophically grown TIE-1 cells were transferred into anoxic [purged with 34.5 kPa N_2_/CO_2_ (80%/20%)] bicarbonate buffered freshwater (FW) (79) medium with ammonium chloride (NH_4_Cl) (5.61 mM) as a nitrogen source and H_2_ as an electron donor. The cells were incubated at 30 °C under a 60-Watt incandescent light source placed at a distance of 12.5 cm. For photoautotrophic growth with Fe(II) as an electron donor, photoautotrophically grown TIE-1 cells were inoculated into anoxic FW medium prepared under the flow of 34.5 kPa N_2_/CO_2_ (80%/20%) and dispensed into pre-sterilized serum bottles/Balch tubes purged with 34.5 kPa N_2_/CO_2_ (80%/20%). The containers were then sealed using sterile butyl rubber stoppers with aluminum crimp and stored at room temperature for at least a day before supplementing with anoxic sterile stocks of FeCl_2_ to a final concentration of 5 mM. For anoxic growth, all sample manipulations were performed inside an anaerobic chamber (Coy, Michigan, USA). For growth on solid medium, LB or YP medium was solidified with 1.5% agar and supplemented with antibiotics as done previously (64).

### DNA isolation, plasmid and strain construction

Genomic DNA of TIE-1 was isolated using the DNeasy Blood and Tissue kit (Qiagen, Valencia, CA, USA) and used as a template for PCR reactions. All nucleic acids isolated in this study were quantified using a Nanodrop 1000 Spectrophotometer (Thermo Scientific, Waltham, MA, USA). A QIAprep Spin Miniprep kit (Qiagen, Valencia, CA, USA) was used for plasmid DNA isolation from *E. coli*. All primers used in this study were obtained from the Integrated DNA Technologies, Coralville, IA, USA, and their sequences are available upon request. The identity of all DNA constructs was confirmed via DNA sequencing at Genewiz Inc., South Plainfield, NJ, USA. *E. coli* strains were transformed by electroporation using an *E. coli* Gene Pulser**®** (Biorad, Hercules, CA, USA), as recommended by the supplier. Plasmids were mobilized from *E. coli* S17-1/λ pir into TIE-1 by conjugation on YP agar. All plasmids constructed are indicated in Table S5. Mutant construction was performed as described previously for TIE-1 (28, 36, 64, 65). Briefly, 1-kb DNA fragment upstream and downstream of the gene of interest were PCR amplified using TIE-1 genomic DNA as the template and cloned into pJQ200KS, a suicide vector. The constructs were mobilized into TIE-1 via conjugation with *E. coli* S17-1. Homologous recombination was allowed to occur on selective YP medium containing 400 µg/ml gentamicin. Vector loss was selected using media containing 10% sucrose. Mutants lacking the target genes were screened using PCR. The mutants created are mentioned in Table S4. The primers used for verification are provided in the Table S7.

### *E. coli* β-galactosidase assays

*E. coli* H1771 derivatives were grown in LB medium at pH 7.0 at 37 °C with shaking at 225 rpm. Four different growth conditions were tested for each strain, **(i)** Fe(III)-replete [50 µM Fe(III)-citrate]; **(ii)** Fe(III)-deplete [50 µM Fe(III)-citrate with 100 µM DiP]; **(iii)** Fe(II)-replete [50 µM ascorbate-reduced Fe(III)-citrate]; and **(iv)** Fe(II)-deplete [50 µM ascorbate-reduced Fe(III)-citrate with 100 µM DiP]. Ascorbate-reduced Fe(III)-citrate was prepared as previously described (32). The *E. Coli* strains were grown with 100 μg/ml of ampicillin to maintain the pWKS30 plasmid derivatives. Overnight cultures of these strains were inoculated into the respective media with a dilution factor of 1:100. The cultures were grown at 37 °C with shaking at 225 rpm for 5 hours. These cells were then assayed for β–galactosidase activity as previously described using the Miller assay (80). β-galactosidase activity expressed in Miller units was calculated using the following formula:

Miller units = (OD_420_ * 1000) / (OD_600_ * V * t), where

V = volume of culture processed for each sample, and t = time elapsed from start to end of the reaction.

### *E. coli* growth curves

*E. coli* GR536 derivatives were maintained in LB medium with pH 7.0 with 25 μg/ml of chloramphenicol and 50 μg/ml of kanamycin. The strains carrying pWKS30 derivatives had 100 μg/ml of ampicillin in addition to chloramphenicol and kanamycin. Overnight cultures of these strains were grown and inoculated into Tris-mineral salts medium with 3 g/L casamino acids and 0.2% sodium gluconate (50 mM Tris-Cl pH 7.0, 80 mM NaCl, 20 mM KCl, 20 mM NH_4_Cl, 1 mM MgCl_2_•6H_2_O, 0.2 mM CaCl_2_•2H_2_O, 5 μM iron ammonium citrate, 3 mM Na_2_SO_4_•10H_2_O, 50 μM Na_2_HPO_4_•12H_2_O) with 0.1X SL6 trace elements solution. An increasing concentration of 2,2’-dipyridyl was added to the medium with the highest concentration at 150 μM. Similar to *E. coli* GR536 derivatives, *E. coli* GG48 derivatives were also maintained in LB medium with pH 7.0 with 25 μg/ml of chloramphenicol and 50 μg/ml of kanamycin. The zinc, cadmium, and copper toxicity tests were performed in LB medium at pH 7.0 with increasing concentrations of added ZnCl_2_ from 0-400 μM, CdCl_2_ from 0-100 μM and CuCl_2_ from 0-400 μM, respectively. The cultures were grown at 37 °C with shaking at 225 rpm. Samples were withdrawn periodically for OD_600_ measurements. *E. Coli ΔcopA ΔcueO ΔcusCFBA:cat* derivatives were maintained in pH 7.0 LB medium with 25 μg/ml chloramphenicol. The strains carrying pWKS30 derivatives had 100 μg/ml ampicillin in addition to chloramphenicol.

### Iron and manganese uptake assays

Iron and manganese uptake assays were performed as previously described with some modifications (48). For anoxic growth conditions, TIE-1 strains were grown in FW medium at pH 7.0 with 1 mM sodium succinate at 30 °C under a 60-Watt incandescent light source placed at a distance of 12.5 cm from the cultures. Time-course OD_660_ was measured by withdrawing the samples under anoxic conditions. Late exponential cultures were then aliquoted into pre-sterilized anoxic Balch tubes followed by head-space flushing with 34.5 kPa N_2_: CO_2_ (80%:20%). The tubes were placed on ice until use. For oxic growth conditions, TIE-1 was grown in 3 g/L yeast extract and 3 g/L peptone media supplemented with 1 mM succinate (YP) with shaking at 250 rpm for 120 hours.

To test iron and manganese accumulation by single-gene mutants, complemented and WT strains and their response to metal-replete and -deplete conditions with respect to oxygen availability, the following experiments were included, **(i)** normal level of Fe(II) or Mn(II) (basal media only); **(ii)** basal media with 100 µM 2,2’-dipyridyl (DiP) or ethylenediaminetetraacetic acid (EDTA); **(iii)** basal media with 50 µM Fe(II) or Mn(II); and **(iv)** basal media with 100 µM of DiP or EDTA and 50 µM Fe(II) or Mn(II), under oxic and anoxic conditions. Fe(II), Mn(II), EDTA, and DP were added from a 50 mM stock solution at the beginning of the growth.

### Iron and manganese analysis by ICP-MS analysis

1 mL sample was taken from the cultures incubated with iron or manganese and centrifuged at 21,000 g for five minutes. Samples were washed with 1mL TE (Tris-ethylene diamine tetra acetic acid) buffer and 1mL of ultrapure water. Washed pellets were then added with 50 µL high purity HNO_3_ trace element (Fisher) and digested with CEM MARS 6 microwave at 180 °C for 20 minutes. Digested samples were then diluted two-fold with ultrapure water and filtered with a 0.22 µm PES filter. Filtered samples were analyzed using Perkin Elmer Elan DRC II ICP-MS. Iron was analyzed as 55.845 isotope and Manganese as 54.94 isotope. Yttrium was used as an internal standard. Standard curves were built from ICP-MS grade Iron and Manganese (Inorganic Venture, VA, USA) using the following concentration: 0, 1, 10, 50, 100 and 200 ppb. Standards were also used as abiotic controls. Obtained Fe and Mn concentrations were normalized to the OD of the sample and further normalized to the concentration obtained from the WT – TIE-1.

### Gene expression analysis

Gene expression analysis was performed using reverse transcription quantitative polymerase chain reaction (RT-qPCR). We quantified mRNA abundance of the putative Fe(II) transporter genes in TIE-1 under, **(i)** basal medium only (Fe ∼4 nM, Mn ∼0.25 nM); **(ii)** basal medium with 100 µM DiP or EDTA; **(iii)** basal media with 50 µM Fe(II) or Mn(II) and; **(iv)** basal medium with 100 µM of DiP or EDTA and 50 µM Fe(II) or Mn(II), under oxic and anoxic conditions. The data were normalized to the gene expression obtained from the basal medium only. The comparative Ct method was used as described previously to assess the expression of the relevant genes. Primer efficiencies were determined using the manufacturer’s protocol (Applied Biosystems Inc. User Bulletin #2). *clpX* and *recA* were used as the two internal standards, which have been previously used and validated as internal standards (64). The primers used for the assays are indicated in Table S6. The iScript cDNA synthesis kit was used for reverse transcription (Biorad, Hercules, CA, USA). iTaq FAST SYBR Green Supermix with ROX (Biorad, Hercules, CA, USA) and the Stratagene Mx3005P QPCR System (Agilent, Santa Clara, CA, USA) were used for all quantitative assays.

### Bioinformatics tools

For identifying iron transporters in TIE-1, delta-blast (81), FASTA (82) (http://www.ebi.ac.uk/Tools/sss/fasta/), and the IMG ortholog neighborhood search were used (83) (http://img.jgi.doe.gov/cgi-bin/w/main.cgi). Transmembrane predictions were made using the TMHMM Server v. 2.0 at http://www.cbs.dtu.dk/services/TMHMM/. Promoter predictions were made using Virtual Footprint v. 3.0 at http://www.prodoric.de/vfp/ (84).

### Statistical analysis

All statistical analyses (two tails Student’s t-test) were performed with Python. *P*-value <0.05 was considered to be significant. All the experiments were carried out using biological triplicates.

## ACKNOWLEDGMENTS

We would like to thank The Nano Research and Environmental Laboratory (NREF) at Washington University in St. Louis for their support to measure iron and manganese using ICP-MS. This work was supported by the following grants to A.B.: The David and Lucile Packard Foundation Fellowship (201563111), the U.S. Department of Energy (grant number DESC0014613), and the U.S. Department of Defense, Army Research Office (grant number W911NF-18-1-0037). A.B. was also funded by a Collaboration Initiation Grant, an Office of the Vice-Chancellor of Research Grant, and an International Center for Energy, Environment and Sustainability Grant from Washington University in St. Louis.

